# Harnessing antimicrobial peptide genes to expedite disease-resistant enhancement in aquaculture: Transgenesis and genome editing

**DOI:** 10.1101/2023.01.05.522886

**Authors:** Jinhai Wang, Yu Cheng

## Abstract

Numerous studies have demonstrated that genome editing and transgenesis by integrating vector-engineered antimicrobial peptide genes (AMGs) is effective to modulate the fish’s innate immune system. To generalize the knowledge of AMG application in aquaculture, here we recruited 540 data entries from a pool of empirical studies, which included 18 peer-reviewed publications and spanning 12 diseases. We systematically re-processed and re-analyzed these data by harnessing a cross-disease meta-analysis. On aggregate, AMG-genetic engineering aimed at enhancing disease resistance was shown to decrease the number of colony-forming units of bacteria, improve lysozyme activity, increase the post-infection survival rates, and alter the expression of AMGs and immune-related genes in aquatic animals. Furthermore, the AMG-pathogen combating activity was triggered within two hours after infection and lasted 48 hours, and the overexpression of AMGs was dominant in the spleen and skin, followed by the kidney and liver during this period. Typically, regardless of the type of AMGs, the synergistic expression of AMGs with IL, IKβ, TGFβ, C3b and TLR in AMG-integrated fish contributed to activating inflammatory/immune responses against pathogens. In addition, innovative CRISPR/Cas9-mediated systems enabling the site-directed knock-in of foreign genes at multiple loci were presented and prospected for disease-resistant enhancement in combination with other favorable fish-producing traits, including fast-growing, sterility, and enriched fatty acid. Altogether, our findings indicated that AMGs as transgenes have substantial potential to modulate the fish’s innate immune system and accelerate disease-resistant enhancement combined with genetic engineering.

## 1. Introduction

Aquaculture plays a critical role in providing essential protein and nutrients to satisfy global food demand for billions of people, as well as employment and other livelihoods. In 2019, aquaculture produced 85.3 million tons of fish, and annual fish and fish product consumption is predicted to reach 25.5 kg per capita by 2050 (FAO, 2022). Such growth would make aquatic food contribute a larger proportion to the food basket globally, assisting in the filling of the food gap. Despite aquaculture being the fastest-growing source of food in the world, sustainable development is under constant threat as infectious diseases afflict cultured aquatic animal populations.

Like other animal-rearing systems, fish farming is plagued by disease issues because of its intensification and commercialization (FAO, 2021). Moreover, farm-raised fish are susceptible to a variety of pathogen infections. On the one hand, strict biosecurity protocols and improved aquaculture practices have been implemented to protect against devastating losses from infectious diseases and make aquaculture more sustainable and environmental-friendly (USDA, 2021). On the other hand, for a long time, the prudent use of veterinary medicines has been one of the most effective strategies to control disease in aquaculture. However, indiscriminate use of these veterinary drugs is contributing to the spread of antimicrobial resistance (AMR), a major concern for the environment and even humans (Bondad-Reantaso et al., 2020; Karunasagar et al., 2020). Although Food and Drug Administration (FDA) has approved the use of three drugs in aquafeeds, including florfenicol, oxytetracycline dihydrate, sulfadimethoxine/ormetoprim and sulfamerazine (FDA, 2022), these approved medications are region- and disease-specific. Recently, World Health Organization (WHO) and Food and Agriculture Organization (FAO) have restricted the use of antimicrobials/antibiotics and veterinary drugs in food-producing animals, including cultured fish (WHO, 2017; FAO, 2021).

In addition to management and regulation by the government, advancements in disease enhancement for cultured aquatic animals include the use of vaccinations (Dadar et al., 2017; Assefa and Abunna, 2018), functional feed additives (Rico et al., 2013; Dawood et al., 2018), and genomic approaches (Houston et al., 2020; Wang et al., 2022a). These advancements not only offer the possibility of reducing or eliminating antibiotic use in the future, but they also have the dual benefits of reducing AMR’s effects while bolstering production. Notably, a considerable number of natural molecules including antimicrobial peptides (AMPs), probiotics, immunostimulants, organic acids and plant extracts are being extensively investigated in aquaculture as therapeutic alternatives to conventional antibiotics (Dawood et al., 2018; Ahmadifar et al., 2021; Wang et al., 2022b).

AMPs are one of the most promising substitutes for traditional antibiotics because of their broad-spectrum antibacterial properties without causing AMR (Hancock et al., 2016; Mookherjee et al., 2020). They have been demonstrated to be efficient not only in the prevention and treatment of human diseases but also in the control of animals’ pathogen infections (Rodrigues et al., 2021; Silveira et al., 2021). Currently, AMPs can be formulated as supplements into aquafeeds, or alternatively, their genes (antimicrobial peptide genes, AMGs) can be integrated into the genomes of fish from one species to another via transgenesis or genome editing, enabling them to be overexpressed when pathogens invade, thereby improving disease resistance (Lo et al., 2014; Wang et al., 2022a). Dunham et al. (2002) first demonstrated that cecropin-transgenic channel catfish (*Ictalurus punctatus*) exhibited strong bactericidal activities against *Flavobacterium columnare* and *Edwardsiella ictaluri* by increasing survival rates compared to the wild-type individuals. Over the following two decades, more investigations revealed that this acquisition of disease resistance was long-lasting and passed down to succeeding generations (Mao et al., 2004; Yazawa et al., 2006; Hsieh et al., 2010; Chiou et al., 2014; Elaswad et al., 2019).

Although the most intuitive advantage of AMG-integrated fish is a significantly higher survival rate when they are infected by pathogens. An increasing number of studies indicated that underlying factors like decreased colony-forming unit (CFU) of bacteria, enhanced enzyme activity, increased expressions of AMGs and immune-related genes contributed to improving the survival rate in individuals harboring foreign AMGs (Pridgeon et al., 2013ab; Lo et al., 2014; Simora et al., 2020). For example, tail tissue of hepcidin-transgenic zebrafish (*Danio rerio*) and convict cichlid (*Archocentrus nigrofasciatus*) exhibited significant reduction in CFU against *Vibrio vulnificus* compared to wild-type fish (Hsieh et al., 2010). Besides, the caudal peduncle from transgenic zebrafish carrying epinecidin inhibited the bacterial growth when fish were challenged with *V. vulnificus* (Peng et al., 2010). Pridgeon et al. (2013b) have proven that recombinant goose-type lysozyme enhanced innate lysozyme activity in the serum of channel catfish after *Micrococcus lysodeikticus* infection. In addition, the expression of granulin peptide (GRN-41) was induced in the spleen of GRN-41-transgenic zebrafish at 6 hours post *V. vulnificus* infection (Wu et al., 2018). Interestingly, cathelicidin was significantly overexpressed in variety of tissues in cathelicidin-integrated channel catfish even in the absence of pathogenic infections (Simora et al., 2020). Furthermore, a considerable number of innate immune-related genes can be induced or activated in AMG-integrated fish after pathogen infections. Correspondingly, Lo et al. (2014) discovered that more than 2,000 immune-relevant genes were differentially expressed in the spleen, liver and kidney of the cecropin-transgenic rainbow trout (*Oncorhynchus mykiss*) when compared to non-transgenic fish. More specifically, after infection with *V. vulnificus*, both hepcidin- and epinecidin-transgenic zebrafish displayed elevated expressions of myeloid differentiation primary response gene (Myd88), interleukin (IL)-10, IL-26 and toll-like receptor (TLR)-4a, but IL-1β and IL-15 were downregulated (Hsieh et al., 2010; Peng et al., 2010). In aggregate, in addition to directly killing pathogens, AMG-integrated fish also have a variety of immunomodulatory effects to boost host defenses.

Numerous investigations revealed that foreign AMGs as transgenes always bring positive effects in killing pathogens, boosting the innate immune system and improving disease resistance in aquatic animals. Nevertheless, various and even incompatible conclusions emerged due to the variety of the independent studies. Herein, compared to non-transgenic individuals, goose lysozyme significantly enhanced innate lysozyme activity in channel catfish after bacterial infection (Pridgeon et al., 2013b). Contrarily, serum lysozyme activity showed no significant differences between the lactoferrin-transgenic and control groups when grass carp (*Ctenopharyngodon idellus*) were injected with *Aeromonas hydrophila* (Mao et al., 2004). Similarly, the inhibitory effect against *V. vulnificus* was greatly improved when tilapia hepcidin was integrated into the zebrafish genome, but not against *Streptococcus agalactiae* (Hsieh et al., 2010). Furthermore, Wu et al. (2018) stated that tilapia GRN-41 can elevate the expression of IL-1β after the *V. vulnificus* challenge in transgenic zebrafish. However, following the same bacterial infection, IL-1β was downregulated in hepcidin- and epinecidin-transgenic zebrafish (Hsieh et al., 2010; Peng et al., 2010). In this instance, if credible conclusions can be drawn, it is very crucial to statistically assess and synthesize the existing findings in addition to undertaking more investigations to gather data.

Several variables, including the type of AMGs, fish species, and pathogen type, are identified by individual studies as having an impact on the outcomes of disease-resistant improvement. As a result, the estimated results of increased disease resistance are therefore heterogeneous due to the specificity of these factors, and this high variability needs to be considered when assessing the effect of AMG application in aquaculture. In this study, we delved deeper into how these general determinants influence disease-resistant enhancement through multiple moderator analyses based on our initial meta-data. Therefore, the main objective of this study is to quantitatively integrate empirical data on the use of AMGs through transgenesis or genome editing in aquaculture and aim to 1) Identify consistencies across studies for AMG application to assess the performance of AMG-integrated fish in terms of bacterial CFU, lysozyme activity (LYA), cumulative survival rate (CSR), and the expression of exogenous AMGs (TGE) and immune-related genes (IRGE) after an invasion of pathogens. 2) Confirm whether the pathogen-combating abilities of AMGs are fish- or disease-specific. 3) Determine if there are any key immune-related genes that collaborate with exogenous AMGs to trigger the innate immune system. 4) Identify tissue distribution and temporal patterns of these genes’ expression. Herein, conducting a cross-disease meta-analysis based on the global synthesis of published data, we are able to generalize how to engineer AMGs to boost disease enhancement in aquaculture through genetic engineering.

## 2. Data compilation and analysis

The current meta-analysis followed the Preferred Reporting Items for Systematic Reviews and Meta-Analyses (PRISMA) guideline (Moher et al., 2009) in literature search, study selection, and data collection.

### 2.1. Literature search

The PRISMA flowchart demonstrated the search procedure (Fig. S1). The search and collection of literature were carried out on multiple databases such as Web of Science (WOS), PubMed, Aquatic Sciences and Fisheries Abstracts (ASFA) and Academic Search Premier (ASP) using keywords (“transgen*” OR “genome editing” OR “gene editing” OR “CRISPR/Cas9” OR “microinjection” OR “electroporation”) AND (“antimicrobial peptide” OR “AMP” OR “antimicrobial peptide gene” OR “AMG”) AND (“aquaculture” OR “aquatic animals” OR “fish” OR “marine” OR “shellfish” OR “shrimp”) AND (“disease resistance” OR “bactericidal” OR “antibacterial” OR “antiviral” OR “antiparasitic” OR “phagocytic activity” OR “bacterial activity” OR “immune response”), and the language was limited to English. Furthermore, relevant literature from empirical collections was chosen and grouped as “additional records”. The last retrieval date was 5/4/2022 for these online databases. Initially, 115 articles were gathered using the keywords mentioned above. In addition to peer-reviewed scientific articles, some experimentally acquired but unpublished data from our team was included in the current meta-analysis. Following a thorough review of abstracts and full texts, 18 publications and 3 unpublished papers (from our team) comprising a total of 540 data entries were adopted to build the database as shown in Supplementary S1.

### 2.2. Selection criteria

The following criteria were included: 1) Application scope and description of antimicrobial peptide genes (AMGs). Types, sequences or thorough descriptions of AMGs are revealed in the retrieved publications, and foreign AMGs must be applied as transgenes by harnessing transgenesis or genome editing for fish, shellfish or shrimp. 2) Pathogen species. The pathogens’ names or descriptions should be represented if collected publications employed pathogen-challenge experiments. 3) Complete available data. Recruited articles should include CFU of pathogens, LYA, CSR, TGE or IRGE after pathogen infection. All these parameters should be present in both control (non-edited) and transgenic/gene-edited (AMG-integrated) groups. For that literature that fails to report standard deviations (SDs) or standard errors (SEs) and cannot be inferred from existing data, we automatically fill in by the imputation method (Furukawa et al., 2006). For publications by Dunham et al. (2002), Sarmasik et al. (2002), and Mao et al. (2004), this generic strategy was employed.

The exclusion criteria were listed below: 1) AMGs are not employed directly as transgenes in aquacultured species. For instance, the study by Lin et al. (2010) revealed that disease resistance of zebrafish (*Danio rerio*) against multiple bacteria improved when fed with lactoferricin-transgenic fish embryos, which was not within our scope to investigate. 2) Uncertified AMGs. Although the myxovirus resistance (Mx) gene is extremely anti-virus and has been used for disease enhancement against grass carp reovirus for a rare minnow (*Gobiocypris rarus*) as a transgene (Su et al., 2009), no documentation has classed it as an AMG. In this case, we excluded it in our meta-analysis. 3) Data is incomplete. Lo et al. (2014) and Han et al. (2018) determined extensive gene expression patterns in cecropin P1 transgenic rainbow trout via microarray and RNA sequencing after pathogen infection, but we could not extract it due to a lack of specific gene expression data.

### 2.3. Information extraction

The following information was extracted from each selected study: author information (first author, year), article title, AMG type, pathogen type, fish species and sample size, as well as mean values and SDs of outcome data in non-edited and AMG-integrated groups, including CFU of bacteria, LYA, CSR, TGE or IRGE after pathogen infection. In addition, if just SE and sample size (n) are supplied in a single publication, SD should be determined using the formula SD = SE*(n)^1/2^. We used ImageJ to extract data (mean and SD or SE) from articles that only provided figures (https://imagej.nih.gov/ij/).

### 2.4. Statistical analysis

All statistical analyses were conducted using “metafor” and “orchaRd” packages in RStudio version 3.6.3 (2020-02-29). The significance level for all statistical tests was set to *P* < 0.05. R codes were attached in Supplementary S2.

#### 2.4.1. Effect size calculations

The goal was to compare the disease resistance of AMG-integrated fish to wild-type individuals using the CFU, LYA, CSR, TGE, and IRGE parameters. Here, these five outcomes are continuous variables, and the sample sizes are not identical for non-edited and AMG-integrated groups in our cases. As a result, Hedges’*g* was employed to calculate effect sizes for our meta-analysis (Hedges and Olkin, 2014). Using random-effects models, we estimated the overall mean effect sizes with 95% confidence intervals (CIs) and 95% prediction intervals (PIs) to compare the effects of non-edited and AMG-integrated groups for each parameter. If the 95% CIs do not intersect with zero, they are considerably different between these two groups in statistics. Positive values of mean effect sizes for LYA, CSR, TGE and IRGE indicate that the AMGs improve disease resistance through altering these parameters compared with the non-edited group. On the contrary, negative values indicate that the performance declined with the presence of AMG in transgenic fish. Negative effect size for CFU shows that AMG transgenic fish have increased bacteriostatic activity by lowering colony-forming units. Furthermore, Hedges’*g* is interpreted using a rule of thumb similar to Cohen’s *d* but with some modifications: 0 < | Hedges’*g* | ≤ 0.5, small effect; 0.5 < | Hedges’*g* | ≤ 0.8, medium effect and | Hedges’*g* | > 0.8, large effect (Cohen, 1977). For example, an effect size less than 0.5 would likely be considered a weak effect. It means that even if the difference between the two groups is statistically significant (*P* < 0.05), the actual difference between the group means is trivial. The *I*^2^ statistic (Higgins and Thompson, 2002) was used to calculate the percent variance owing to inconsistencies in the population effect across studies, and *I*^2^ > 50% indicated significant between-study heterogeneity. In the analysis that follows, moderator tests were performed.

#### 2.4.2. Publication bias and sensitivity analysis

To assess publication bias, funnel plots were created, and all estimates were subjected to a classic Egger’s regression test for funnel plot asymmetry. A *P* value < 0.05 implies that the funnel plot is asymmetric, and publication bias is possible. In this case, Tweedie’s nonparametric “trim and fill” approach (Duval and Tweedie, 2000) should be used to see if random-generated studies are needed to reduce potential publication bias.

We identified outliers and influential observations of the estimated measures across studies using Cook’s distance, and the influence and leave-one-out diagnostics (Viechtbauer and Cheung, 2010) as a combined strategy to evaluate the stability and reliability of our meta-analysis. After detecting the significant outliers by sensitivity analysis, we recalculated the pooled effect sizes by excluding the outlier observations to assess the final effect without outliers.

#### 2.4.3. Moderator analysis

In principle, an *I*^2^ > 50% indicates significant heterogeneity across studies and they need to be regrouped for further moderator analyses using mixed-effect models. By combining the pool of empirical studies, we recognized that fish species, AMG type and pathogen type have decisive effects on primary response variables CFU, LYA and CSR. Therefore, fish-, AMG- and pathogen-moderator analyses were performed for these three parameters. With respect to LYA, all the studies were focused on disease-resistant enhancement against *A. hydrophila*, thus we only tested fish and AMG moderators. To increase the sample size of effect sizes, we regrouped homologous AMGs into broader categories (e.g., cecropin B and cecropin P1 belonged to cecropin; chicken lysozyme and goose lysozyme were grouped into lysozyme) for AMG-moderator analysis of CSR. Additionally, the same strategy was also adopted in the pathogen-moderator analysis. For instance, *Edwardsiella ictaluri* and *E. tarda* were broadened into *Edwardsiella*, while *Aeromonas salmonicida* and *A. hydrophila* were categorized into *Aeromonas*.

Gene expression is highly specific after exogenous AMGs are integrated into the genomes of targeted species, particularly in tissue and time (Wang et al., 2022a). Consequently, in addition to fish- and AMG-moderator analyses, the tissue-moderator test was employed to determine the specificity of AMG expression in different tissues for TGE. However, we did not conduct the time-moderator analysis for TGE since all the gene expression data was taken at zero time point (0-hour), which indicated it was collected before the fish were infected with pathogens.

Like TGE, immune-related genes also have a variety of expression patterns. Therefore, IRGE was subjected to fish-, gene-, and time-moderator analyses. Currently, the expression level of 26 immune-related genes has been investigated after pathogen infection (Supplementary S1). We also regrouped homologous genes into broader groups to expand the sample size (for example, IL-1β, IL-8, IL-10, IL-15, IL21, IL-22 and IL-26 belonged to the IL gene; TLR4α, TLR1, TLR3 and TLR5 were combined into TLR and so on). In this case, ten subgroups (IL, TLR, TNF, C3b, IKβ, TGFβ, NFKβ, TRAM1, MyD88 and Lysozyme) were set up for the gene-moderator analysis. The expression of immune-related genes has been widely reported at various time points (from 0 to 48 hours). Thus, ten groups spanning a 48-hour interval (0-hour, 1-hour, 2-hour, 3-hour, 4-hour, 6-hour, 8-hour, 12-hour, 24-hour and 48-hour) were selected for the time-moderator analysis.

## 3. Results

The current meta-analysis extracted 74 figures from eight studies, including line charts and histograms (Supplementary S3). The 540 effect sizes (k = 14 for CFU, k = 3 for LYA, k = 50 for SCR, k = 42 for TGE, and k = 431 for IRGE) from 18 studies spanning 8 fish species and 12 diseases were included in the recruited dataset (Dunham et al., 2002; Sarmasik et al., 2002; Zhong et al., 2002; Mao et al., 2004; Yazawa et al., 2006; Lin et al., 2009; Hsieh et al., 2010; Peng et al., 2010; Pan et al., 2011; Chiou et al., 2014; Pridgeon et al., 2013ab; Lee et al., 2013; Lin et al., 2016; Wu et al., 2018; Su et al., 2018; Elaswad et al., 2019; Simora et al., 2020). In total, 15 AMGs were integrated into the fish genome in an attempt to improve disease resistance via genetic engineering (Fig. 1).

**Fig. 1.**
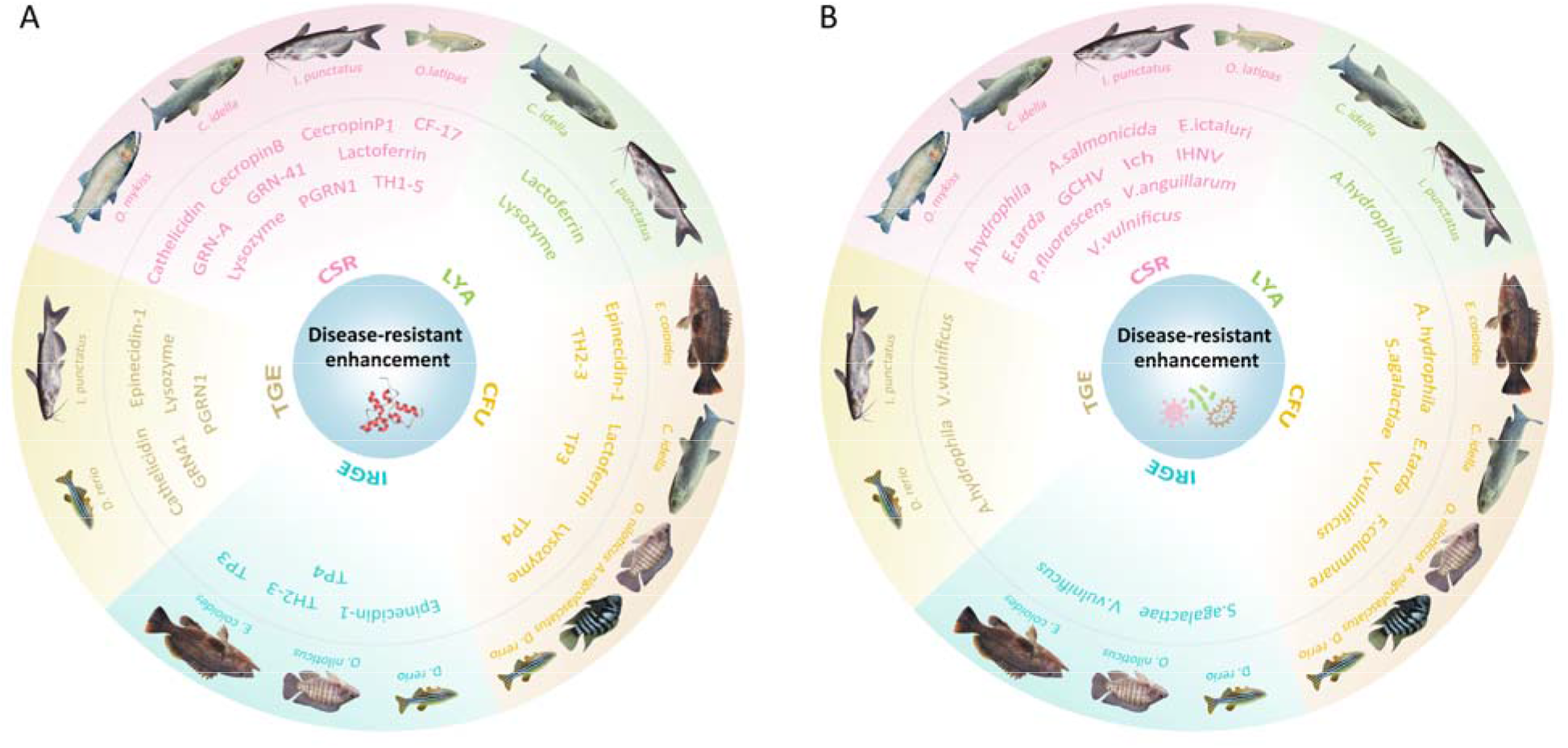
A summary of the current disease-resistant enhancement applications in aquaculture using antimicrobial peptide genes (AMGs) combined with transgenesis and genome editing. This field was covered by 18 articles, which included 8 fish species, 15 AMGs, and 12 diseases. (**A**) Different AMGs were applied for each parameter. For example, two AMGs (lactoferrin and lysozyme) are used as transgenes in grass carp (*C. idella*) and channel catfish (*I. punctatus*) for evaluation of LYA. (**B**) A variety of pathogens were involved for each parameter. For example, one bacterial species (*A. hydrophila*) is used as a pathogenic infection in grass carp (*C. idella*) and channel catfish (*I. punctatus*) for evaluation of LYA. For full scientific names of the fish and pathogens please refer to Supplementary S1. Ich, *Ichthyophthirius multifiliis*; GCHV, grass carp hemorrhage virus; IHNV, infectious hematopoietic necrosis virus; TP3, tilapia piscidin 3; TP4, tilapia piscidin 4; TH2-3, tilapia hepcidin 2-3; TH1-5, tilapia hepcidin 1-5; PGRN1, a type of progranulin gene from Mozambique tilapia; GRN-41/GRN-A, AMGs from Mozambique tilapia to produce secreted GRN peptides; CF-17, a synthetic cecropin B analog; IRGE, the expression of immune-related genes; TGE, the expression of exogenous AMGs; CSR, cumulative survival rate; LYA, lysozyme activity; CFU, colony-forming unit of bacteria.

### 3.1. Overall effect summary

Table 1 summarizes the main findings from our meta-analysis. AMGs showed negative effects on CFU (mean effect = −2.76, *P* < 0.0001, k = 14) based on the overall effect size combining all studies, but positive effects on LYA (mean effect = 3.44, *P* = 0.0815, k = 3), CSR (mean effect = 7.54, *P* < 0.0001, k = 50) (Fig. 2), TGE (mean effect = 4.51, *P* < 0.0001, k = 42), and IRGE (mean effect = 0.55, *P* = 0.3929, k = 431) (Fig. 3). Overall, the effect sizes of these parameters displayed that AMG-integrated individuals had a lower CFU, a higher LYA, an increased CSR, and an elevated TGE and IRGE compared to non-edited fish.

**Table 1.**
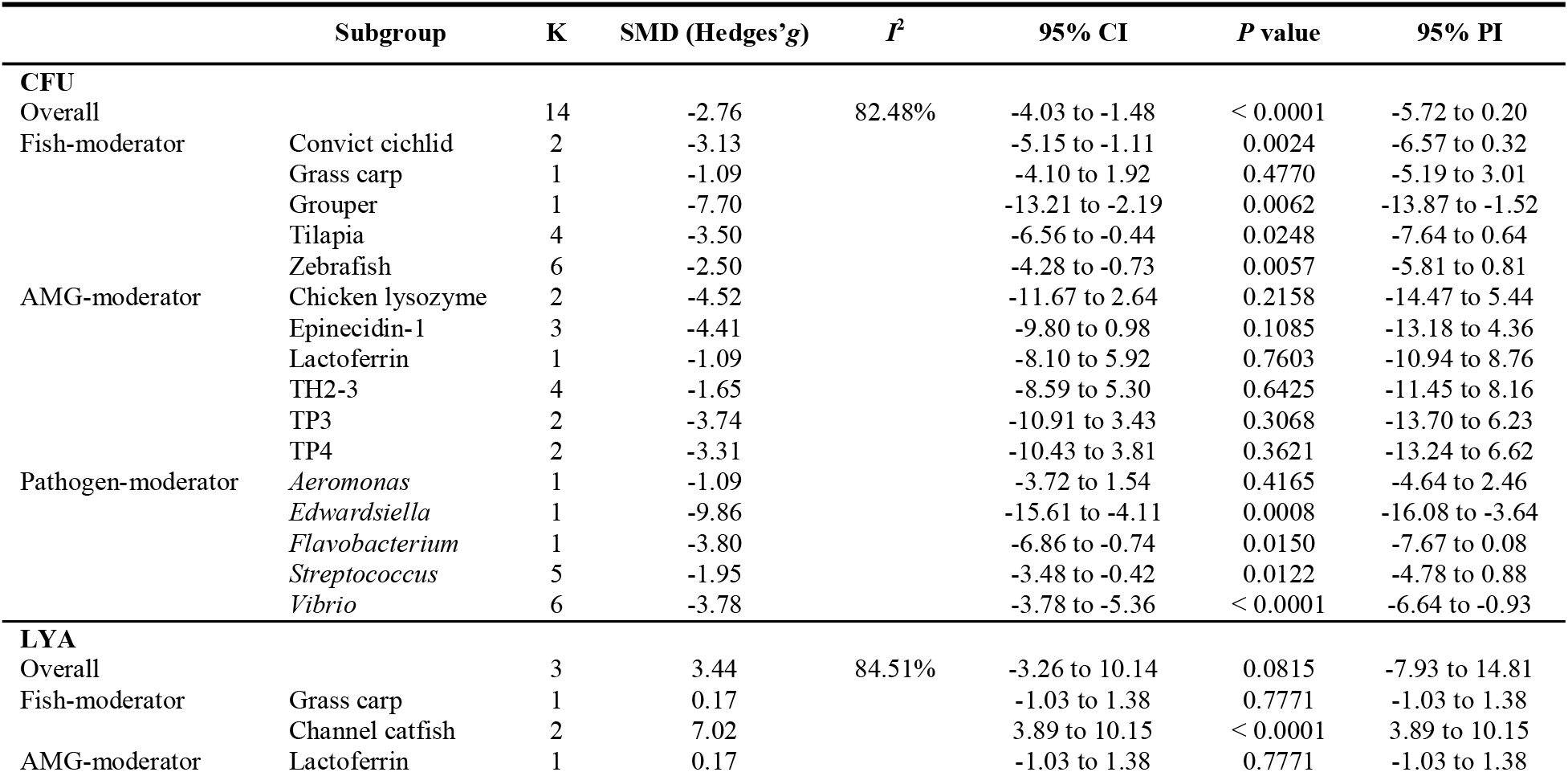
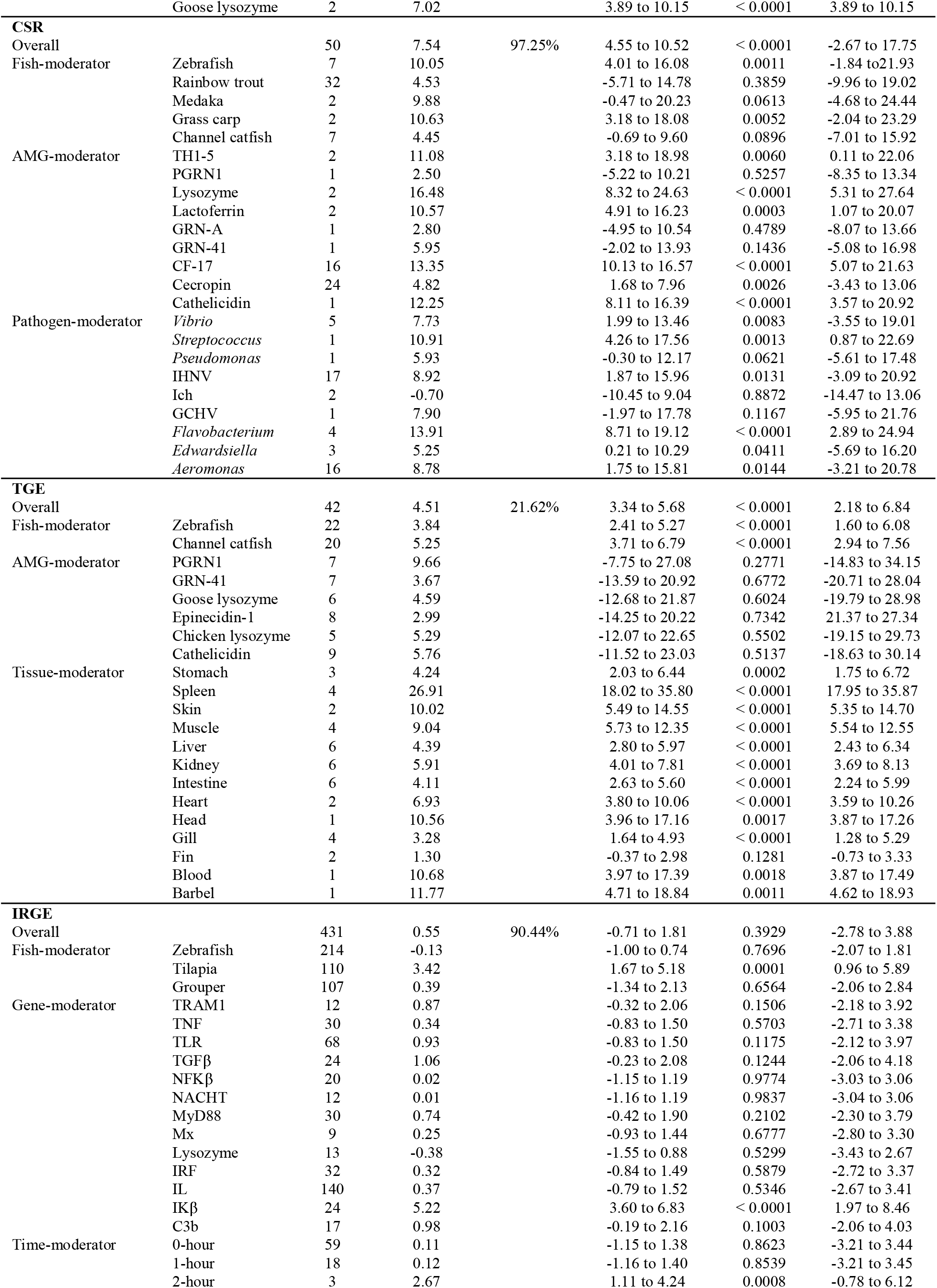
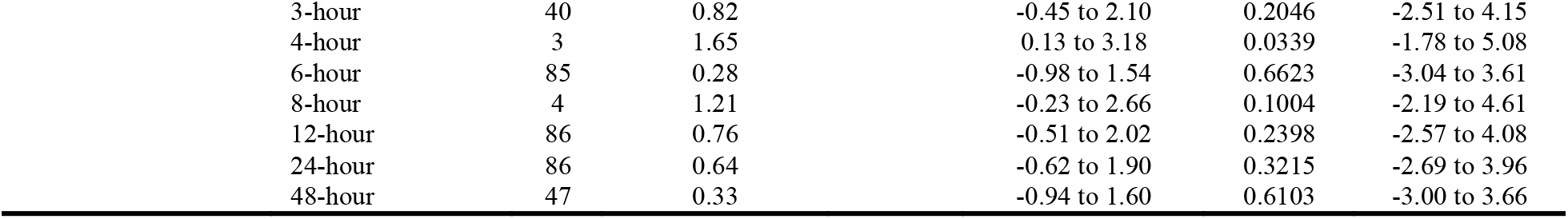
Overall effect size calculations and moderator-analysis results for colony-forming unit (CFU) of bacteria, lysozyme activity (LYA), cumulative survival rate (CSR), the expression of AMGs (TGE) and the expression of immune-related genes (IRGE). The number of effect sizes for each level of these five parameters is indicated by K; SMD, standardized mean difference; *I*^2^, the % variance due to inconsistencies between the population effect across the studies; 95% CI, 95% confidence interval; 95% PI, 95% predicted interval; Ich, *Ichthyophthirius multifiliis*; GCHV, grass carp hemorrhage virus; IHNV, infectious hematopoietic necrosis virus; TP3, tilapia piscidin 3; TP4, tilapia piscidin 4; TH2-3, tilapia hepcidin 2-3; TH1-5, tilapia hepcidin 1-5; PGRN1, a type of progranulin gene from Mozambique tilapia; GRN-41/GRN-A, AMGs from Mozambique tilapia to produce secreted granulin peptides; CF-17, a synthetic cecropin B analog; AMG, antimicrobial peptide gene.

**Fig. 2.**
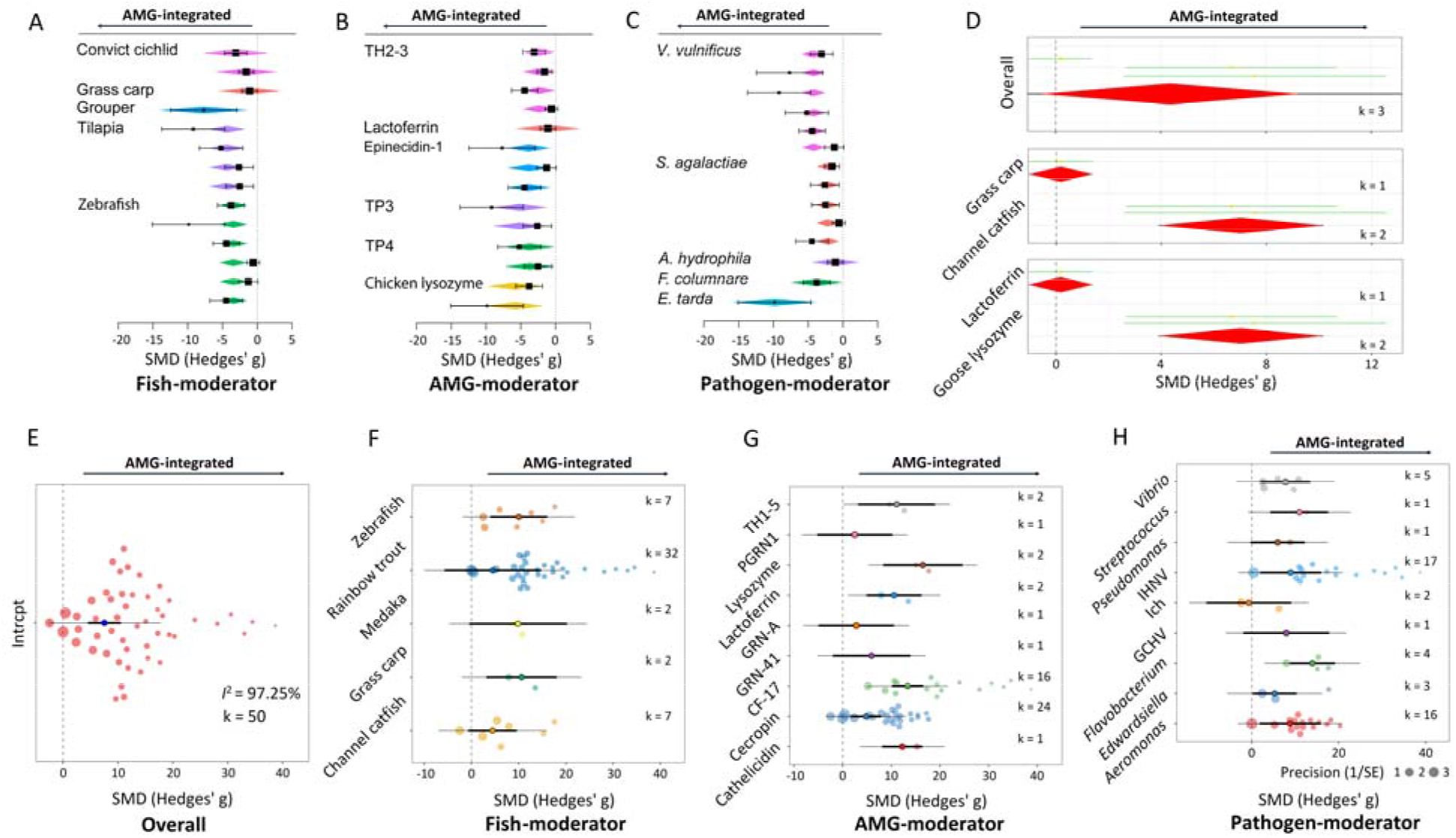
Meta-analytic results of the influence of antimicrobial peptide genes (AMGs) on bacteriostatic action, enzyme activity and survival rate using the standardized mean difference (SMD, Hedges’*g*) as the effect size. Forest plots of integrated AMGs impact on colony-forming unit (CFU). Fish species (**A**), AMG type (**B**) and pathogen type (**C**) were added as moderators, respectively. The black boxplot set of each forest plot depicts the overall effect without any moderators, while the colorful rhombus set illustrates the effect after moderators are added to the model. Categories of each moderator included in this meta-analysis are displayed to the left of the forest plots, and different hues represent distinct categories. (**D**) Caterpillar plot illustrating the effect of integrated AMGs on lysozyme activity (LYA) using Hedges’*g* as the effect size. LYA underwent overall effect and fish/AMG-moderator analyses. Here, the same caterpillar plot displayed that both fish- and AMG-moderator tests yielded the same result. Each green line shows an effect size with a yellow dot (mean effect size). The overall effect size is represented by the red rhombus, which includes a mean value (centre), 95% confidence interval (CI) (left and right borders), and 95% predicted interval (PI) (black line through the rhombus). Orchard plots of integrated AMGs impact on cumulative survival rate (SGR), and overall effect, as well as species-, AMG- and pathogen-moderator analyses (**E - H**) were illustrated. Here are 5, 9 and 9 categories for fish-, AMG- and pathogen-moderator, respectively. K denotes the number of effect sizes for each category of different moderators. For example, k = 7, 32, 2, 2 and 7 indicated that 7, 32, 2, 2 and 7 effect sizes were computed for zebrafish, rainbow trout, medaka, grass carp and channel catfish, respectively, when fish species was considered as a moderator. SMD is represented by 95% CIs and 95% PIs as scaled effect-size points for each study. Each colorful circle shows a scaled different size of the effect. *I*^2^, the percent variance due to inconsistencies across the studies’ population effect. AMG-integrated, gene-edited fish possess an exogenous AMG integrated into the genome via transgenic or genome editing technology.

**Fig. 3.**
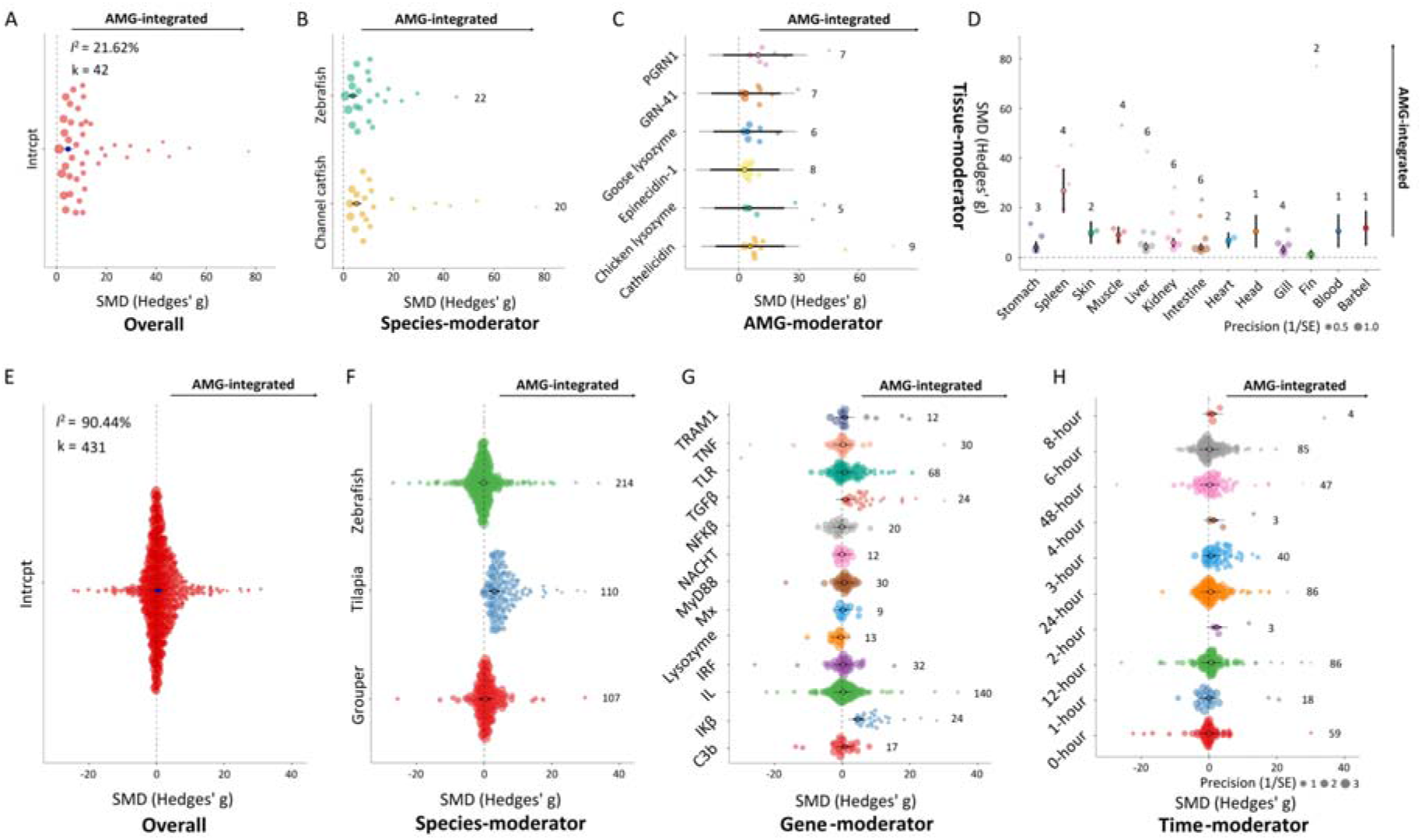
Meta-analytic outcomes of the effect of antimicrobial peptide genes (AMGs) on the level of gene expression, as measured by the standardized mean difference (SMD, Hedges’*g*). Orchard plots of integrated AMGs’ influence on the expression of AMGs (TGE), and overall effect analysis, fish-, AMG- and tissue-moderator analyses (**A - D**) were performed. Here are 2, 6 and 13 categories for fish-, AMG- and tissue-moderator, respectively. Orchard plots of integrated AMGs impact on the expression of immune-related genes (IRGE), and overall effect analysis, fish-, gene- and time-moderator analyses (**E - H**) were conducted. Here are 3, 13 and 10 categories for fish-, gene- and time-moderator, respectively. K indicates the number of effect sizes for each category of different moderators. For example, k = 22 and 20 showed that 22 and 20 effect sizes were estimated for zebrafish and channel catfish, respectively, when fish species was regarded as a moderator. For more specific schematic examples, please see Figure 2.

In addition, the recruited articles represented high heterogeneity in CFU, LYA, CSR and IRGE (*I*^2^ = 82.48% for CFU, *I*^2^ = 84.51% for LYA, *I*^2^ = 97.25% for CSR and *I*^2^ = 90.44% for IRGE) but a low heterogeneity for TGE (*I*^2^ = 21.62%) (Table 1).

### 3.2. Publication bias and sensitivity

Except for LYA (*P* = 0.0578), the funnel plots and Egger’s regression test demonstrated that there was potential publication bias for the effect sizes of CFU, CSR, TGE and IRGE (*P* < 0.0001) (Fig. 4A – E)). In these cases, the “trim and fill” method was used to estimate the number of missing effect sizes or studies in the current meta-analysis for these four parameters. The results revealed that 5, 0, 0, 17 and 97 additional effect sizes/studies were needed to add and balance the publication bias, correspondingly. Specifically, three of the putative five missing effect sizes/studies of CFU were not statistically significant. With respect to TGE, 9 of 17 and additional effect sizes/studies were not statistically significant. However, all 97 missing effect sizes/studies were significant (Fig. 4F – J)). Nevertheless, AMGs still had a significant effect on CFU (adjusted mean effect = −2.23, *P* = 0.0037, k = 19; fail-safe n = 301), TGE (adjusted mean effect = 5.55, *P* < 0.0001, k = 59; fail-safe n = 2257) after incorporating these randomly generated effect sizes/studies. In contrast, incorporation of those 97 created effect sizes/studies reduced the effect on IRGE without a significant difference (adjusted mean effect = 0.43, *P* = 0.1584, k = 528; fail-safe n = 0).

**Fig. 4.**
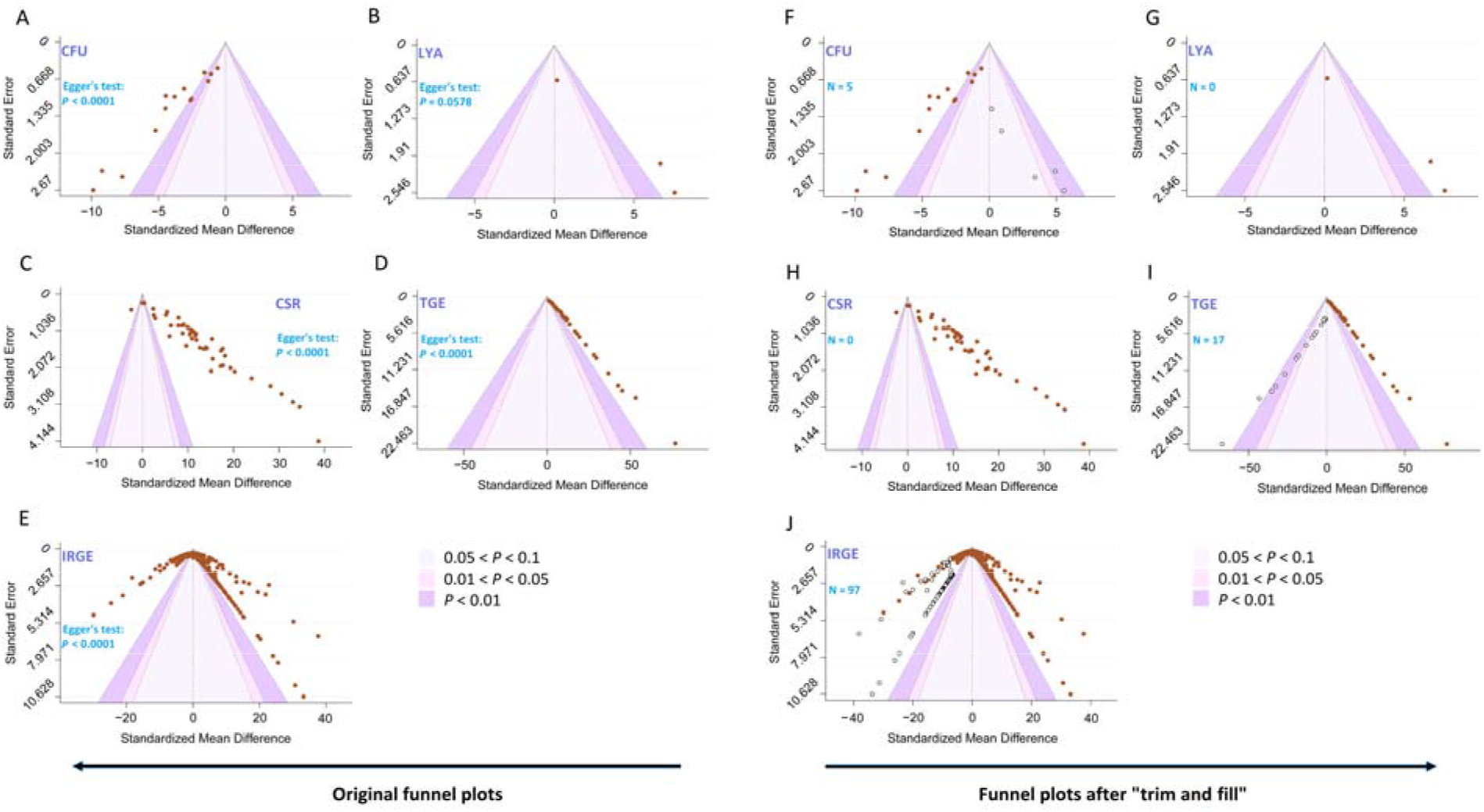
Evidence of publication bias. Contour-enhanced funnel plots of standardized mean difference (Hedges’*g*) for colony-forming unit (CFU) of bacteria, lysozyme activity (LYA), cumulative survival rate (CSR), the expression of AMGs (TGE) and the expression of immune-related genes (IRGE) before and after “trim and fill” analysis. Egger’s regression test, accompanying *P* values and confidence intervals (CIs) were illustrated in each panel. For CFU (**A**), CSR (**C**), TGE (**D**) and IRGE (**E**), funnel plots exhibited substantial asymmetry (*P* < 0.05), indicating potential publication bias. After “trim and fill”, 5, 0, 0, 17 and 97 additional effect sizes/studies need to be implanted to eliminate publication bias for CFU (**F**), LYA (**G**), CSR (**H**), TGE (**I**) and IRGE (**J**), respectively. The reconstructed funnel plots show the additional missing effect sizes imputed in white dots by “trim and fill” if extra studies are needed. 0.05 < *P* < 0.1, 0.01 < *P* < 0.05 and *P* < 0.01 with different colors present 90% CI, 95% CI and 99% CI, respectively. N indicates the number of extra effect sizes/studies that need to be added, and brown dots represent the effect sizes from recruited studies in the current meta-analysis. For example, N = 5 for CFU (**F**) suggests that 5 potential effect sizes/studies need to be added to counteract publication bias. Furthermore, three of them were outside the 90% CI region, implying that these three investigations were not statistically significant.

The influence analysis indicated no potential effect-size outliers for CFU, CSR and TGE, but one and six outliers of effect sizes were represented in LYA and IRGE, respectively (Fig. S2). However, the leave-one-out analysis confirmed that the study from Mao et al. (2004) was an influential case from the overall level for LYA. In the case of IRGE, the result showed that no studies carried a significant deviation of effect size from the overall level if we removed these six unusual cases one by one (Table S1 – S2).

### 3.3. Moderators affect disease-resistant enhancement

Not surprisingly, compared to non-edited fish, there were statistically significant differences in effect sizes of these parameters for AMG-integrated individuals when AMG type, fish species and pathogen type were considered moderators, respectively (Table 1).

#### 3.3.1. Reduced bacterial (CFU)

Antimicrobial peptide genes (AMGs) exhibited large negative effects (all |effect size| > 0.8) on the bacterial CFU even fish species, AMG type and pathogen type were involved in the moderator analyses. Specifically, AMGs had significant negative effects on CFU in convict cichlid, grouper (*Epinephelus coioides*), Nile tilapia (*Oreochromis niloticus*) and zebrafish (*Danio rerio*) (mean effect = −3.13, *P* = 0.0024, k = 2 for convict cichlid; mean effect = −7.70, *P* = 0.0062, k = 1 for grouper; mean effect = −3.50, *P* = 0.0248, k = 4 for tilapia; mean effect = −2.50, *P* = 0.0057, k = 6 for zebrafish), but not in grass carp (mean effect = −1.09, *P* = 0.4770, k = 1) (Fig. 2A). In our example, an effect size of −1.09 would likely be considered a large effect (|mean effect| = 1.09 > 0.8). It means that even if the mean difference between AMG-integrated and control groups is not statistically significant (*P* = 0.4770 > 0.05), the actual difference between the group means is of interest, and the subsequent interpretation of the data also complies with this definition’s standard guideline.

AMG-moderator analysis further revealed that all types of AMGs had large negative effects on CFU. Chicken lysozyme was shown to have the largest effect size, followed by epinecidin-1, TP3, TP4, TH2-3 and lactoferrin (mean effect = −4.52, *P* = 0.2158, k = 2 for chicken lysozyme; mean effect = −4.41, *P* = 0.1085, k = 3 for epinecidin-1; mean effect = −3.74, *P* = 0.3068, k = 2 for TP3; mean effect = −3.31, *P* = 0.3621, k = 2 for TP4; mean effect = −1.65, *P* = 0.6425, k = 4 for TH2-3; mean effect = −1.09, *P* = 0.7603, k = 1 for lactoferrin) (Fig. 2B). Similarly, these various AMGs inhibited the growth of five main bacteria according to the pathogen-moderator test (Fig. 2C).

#### 3.3.2. Increased Lysozyme activity (LYA)

In comparison to the control group, fish with AMG integration have higher lysozyme activity (LYA) with positive effect sizes. According to species-moderator analysis, AMG had a largely beneficial influence on LYA in channel catfish (mean effect = 7.02 > 0.8, *P* < 0.0001, k = 2), which was twice as large as the overall level (mean effect = 3.44). In grass carp, however, this effect was small (mean effect = 0.17 < 0.5, *P* = 0.7771, k = 1). Similar to the results of fish-moderator analysis, goose lysozyme represented a greater effect than lactoferrin on LYA (Fig. 2D).

#### 3.3.3. Improved Cumulative survival rate (CSR)

Antimicrobial peptide genes showed large positive effects on the cumulative survival rate (CSR) after pathogen infections (all Hedges’*g* > 0.8) in all five fish species compared to the control group. Fish-moderator test determined that the largest effect of AMGs on CSR was in grass carp (mean effect = 10.63 > 0.8, *P* = 0.0052, k = 2), followed by zebrafish (mean effect = 10.05 > 0.8, *P* = 0.0011, k = 7), medaka (*Oryzias latipes*) (mean effect = 9.88 > 0.8, *P* = 0.0613, k = 2), rainbow trout (*Oncorhynchus mykiss*) (mean effect = 4.53 > 0.8, *P* = 0.3859, k = 32) and channel catfish (mean effect = 4.45 > 0.8, *P* = 0.0896, k = 2) (Fig. 2F).

The AMG-moderator analysis indicated that different AMGs showed large effects on CSR, with effect sizes ranging from 2.50 to 16.48. Lysozyme, CF-17, cathelicidin, TH1-5, lactoferrin and cecropin showed significant effects on CSR based on the effect size calculations (mean effect = 16.48 > 0.8, *P* < 0.0001, k = 2 for lysozyme; mean effect = 13.35 > 0.8, *P* < 0.0001, k = 16 for CF-17; mean effect = 12.25 > 0.8, *P* < 0.0001, k = 1 for cathelicidin; mean effect = 11.05 > 0.8, *P* = 0.0060, k = 2 for TH1-5; mean effect = 10.57 > 0.8, *P* = 0.0003, k = 2 for lactoferrin; mean effect = 4.82 > 0.8, *P* = 0.0026, k = 24 for cecropin). However, this effect was not significant when PGRN1, GRN-A or GRN-41was integrated into the fish genome (mean effect = 2.50, *P* = 0.5257, k = 1 for PGRN1; mean effect = 2.80, *P* = 0.4789, k = 1 for GRN-A; mean effect = 5.95, *P* = 0.1436, k = 1 for PGRN1) (Fig. 2G).

Furthermore, AMGs demonstrated a variety of differences in effect sizes of CSR for different pathogen infections. AMG-integrated fish exhibited significant improvement on CSR against *Vibrio, Streptococcus*, IHNV, *Flavobacterium, Edwardsiella*, and *Aeromonas* (mean effect = 7.73, *P* = 0.0083, k = 5 for *Vibrio*; mean effect = 10.91, *P* = 0.0013, k = 1 for *Streptococcus*; mean effect = 8.92, *P* = 0.0131, k = 17 for IHNV; mean effect = 13.91, *P* < 0.0001, k = 4 for *Flavobacterium*; mean effect = 5.25, *P* = 0.0411, k = 3 for *Edwardsiella*; mean effect = 8.78, *P* = 0.0144, k = 16 for *Aeromonas*). Despite the fact that AMGs had large favorable effects on CSR against *Pseudomonas* and GCHV, they were both insignificant (mean effect = 5.93, *P* = 0.0621, k = 1 for *Pseudomonas*; mean effect = 7.90, *P* = 0.1167, k = 1 for GCHV). Intriguingly, there was a negative effect on CSR against Ich (mean effect = −0.70, *P* = 0.8872, k = 2) (Fig. 2H).

#### 3.3.4. Induced expression of exogenous AMGs (TGE)

Although fish integrated with AMGs had overall tendency to elevate TGE, moderators often exhibited various effects on TGE. According to the species-moderator test, AMGs showed significant positive effects on TGE in both zebrafish (mean effect = 3.84, *P* < 0.0001, k = 22) and channel catfish (mean effect = 5.25, *P* < 0.0001, k = 20) with large effect sizes (Fig. 3B).

Six AMGs were involved in the AMG-moderator analysis, and the AMG type impacted TGE with large effect sizes ranging from 2.99 to 9.66. PGRN1, cathelicidin, and chicken lysozyme were the top three with the largest effect sizes, followed by goose lysozyme, GRN-41 and epinecidin-1 (mean effect = 9.66, *P* = 0.2771, k = 7 for PGRN1; mean effect = 5.76, *P* = 0.5137, k = 9 for cathelicidin; mean effect = 5.29, *P* = 0.5502, k = 5 for chicken lysozyme; mean effect = 4.59, *P* = 0.6024, k = 6 for goose lysozyme; mean effect = 3.67, *P* = 0.6772, k = 7 for GRN-41; mean effect = 2.99, *P* = 0.7342, k = 8 for epinecidin-1) (Fig. 3C).

The largest effect size of TGE was found in fish spleen (mean effect = 26.91, *P* < 0.0001, k = 4), followed by barbel, blood, head, skin and muscle (mean effect = 11.77, *P* = 0.0011, k = 1 for barbel; mean effect = 10.68, *P* = 0.0018, k = 1 for blood; mean effect = 10.56, *P* = 0.0017, k = 1 for head; mean effect = 10.02, *P* < 0.0001, k = 2 for skin; mean effect = 9.04, *P* < 0.0001, k = 4 for muscle), while heart, kidney, liver, stomach, intestine, gill and fin had smaller effect sizes ranging from 1.30 to 6.93 (mean effect = 6.93, *P* < 0.0001, k = 2 for heart; mean effect = 5.91, *P* < 0.0001, k = 6 for kidney; mean effect = 4.39, *P* < 0.0001, k = 6 for liver; mean effect = 4.24, *P* = 0.0002, k = 3 for stomach; mean effect = 4.11, *P* < 0.0001, k = 6 for intestine; mean effect = 3.28, *P* < 0.0001, k = 4 for gill; mean effect = 1.30, *P* = 0.1281, k = 2 for fin) (Fig. 3D).

#### 3.3.5. Upregulated expression of immune-rated genes (IRGE)

AMGs had positive effects on IRGE in tilapia and grouper (mean effect = 3.42, *P* = 0.0001, k = 110 for tilapia; mean effect = 0.39, *P* = 0.6564, k = 107 for grouper). However, a negative effect was determined in zebrafish (mean effect = −0.13, *P* = 0.7696, k = 214) based on the species-moderator analysis (Fig. 3F).

Like TGE, IRGE showed gene-specificity according to the gene-moderator test. A total of 13 gene categories was analyzed in the current study, and the findings revealed that AMGs had large effect on genes IKβ, TGFβ, C3b, TLR and TRAM1 (mean effect = 5.22 > 0.8, *P* < 0.0001, k = 24 for IKβ; mean effect = 1.06 > 0.8, *P* = 0.1244, k = 24 for TGFβ; mean effect = 0.98 > 0.8, *P* = 0.1003, k = 17 for C3b; mean effect = 0.93 > 0.8, *P* = 0.1175, k = 68 for TLR; mean effect = 0.87 > 0.8, *P* = 0.1506, k = 12 for TRAM1), medium effect on MyD88 (mean effect = 0.74 > 0.5, *P* = 0.2102, k = 30). And small effects were detected on other 7 immune-related genes, including lysozyme, IL, TNF, IRF, Mx, NFKβ and NACHT (|mean effect| = 0.38 < 0.5, *P* = 0.5299, k = 13 for lysozyme; mean effect = 0.37 < 0.5, *P* = 0.5346, k = 140 for IL; mean effect = 0.34 < 0.5, *P* = 0.5703, k = 30 for TNF; mean effect = 0.32 < 0.5, *P* = 0.5879, k = 32 for IRF; mean effect = 0.25 < 0.5, *P* = 0.6777, k = 9 for Mx; mean effect = 0.02 < 0.5, *P* = 0.9774, k = 20 for NFKβ; mean effect = 0.01 < 0.5, *P* = 0.9837, k = 12 for NACHT) (Fig. 3G).

In addition, the time-moderator analysis determined that the effect sizes of IRGE in AMG-integrated fish fluctuated dramatically over time. The effect sizes were small at 0 and 1 hours post pathogen infection (mean effect = 0.11 < 0.5, *P* = 0.8623, k = 59 for 0-hour; mean effect = 0.12 < 0.5, *P* = 0.8539, k = 18 for 1-hour), but increased to 0.76 and 0.64 at 12 and 24 hours, respectively (mean effect = 0.76, *P* = 0.2398, k = 86 for 12-hour; mean effect = 0.64, *P* = 0.3215, k = 86 for 24-hour). This effect size was then reduced to around half at 48 hours (mean effect = 0.33, *P* = 0.6103, k = 47) (Fig. 3H).

## 4. Discussion

Extensive evidence and our current meta-analysis revealed that AMGs were promising and environmental-friendly substitutes for antibiotics, boosting disease-resistant enhancement in fish. Exogenous AMGs that encode AMPs not only directly kill pathogens, but also improve the cumulative survival rate of individuals by intervening in the innate immune and antioxidant systems. In general, successfully integrated AMGs are both efficiently expressed in the presence or absence of a pathogen infection, which tends to trigger the expression of immune-related genes within two hours after infections, and subsequently increase enzymatic activity to boost bacterial clearance. These alterations to the body’s natural anti-infective defenses have sped up the enhancement of disease resistance.

### 4.1. Publication bias and outliers exist but little influence on conclusions

Publication bias is the systematic under- or over-representation of research with specific outcomes in comparison to the total pool of studies conducted, which should be examined to verify the robustness of the meta-analytic outcomes. In our present study, the Egger’s regression test revealed funnel plot asymmetry from the residuals of the overall meta-analytic model using sampling standard error as a predictor (all *P* < 0.05), indicating potential publication bias. However, there were no studies estimated to be missing for LYA and CSR according to the “trim and fill” analysis, therefore, the results remained the same after this modification. In this sense, our results confirmed that publication bias is not the only factor causing asymmetric funnel plots. Indeed, in addition to non-response bias due to uninterested null results, high heterogeneity across studies also contributes to the asymmetry of funnel plots (Rothstein et al., 2005). With respect to CFU, TGE and IRGE, some missing studies are needed to impute to adjust publication bias. After implanting additional studies, we reanalyzed the data and found that consistent conclusions were confirmed by our meta-analysis. Thus, we can therefore draw the conclusion that publication bias did not cause the results to be overestimated or misinterpreted.

High heterogeneity among studies is common, hence it is inevitable that a few of them are outlying or extremely separated from other studies. Therefore, it is required to detect outliers or influential cases using an influence diagnostic for meta-analysis. In our case, one or multiple outliers of effect sizes were determined for LYA and IRGE, respectively. Nonetheless, the leave-one-out diagnosis showed that our conclusions will not be distorted even if we eliminate these exceptional cases one by one. Notably, Turner et al. (2013) claimed that “small-study effects” made it difficult to detect modest effects in the dataset of a meta-analysis that only included several studies. In this study, there were only three effect sizes from two papers for LYA, thus we were unable to determine whether this detection was reliable due to the limited sample size in this situation even if an outlier was observed from our fitted model. However, the analytic results were robust for IRGE even though there were 6 outliers of effect size as the large sample size (324 *vs* 6) can eliminate the unpredictability without altering our findings. Furthermore, effect sizes with a large number in our unbiased meta-analysis yielded accurate effect estimates.

The current study demonstrates a high degree of homogeneity across studies on TGE, indicating that AMGs are overexpressed once they have been integrated into the target genome in all published articles. Due to the absence of exogenous AMGs in wild-type fish compared to AMG-carrying individuals, there is no expression of exogenous AMG in the control group of all studies, which accounts for the high uniformity. This commonality significantly increases the homogeneity among different studies.

### 4.2. Reduced bacterial load favors the enhanced immunity

Bacterial CFU, or the number of viable bacteria or fungal cells, is a valuable macro indicator linked to improved host immunity, and numerous studies have determined the resistance to bacterial infections by counting CFU *in vitro* from specific tissue cultures (Hsieh et al., 2010; Peng et al., 2010; Lin et al., 2016; Simora et al., 2021). Antimicrobial peptides (AMPs) have been recognized as having anti-infective proprieties against invading pathogens *in vivo* and *in vitro* by reducing bacterial load. According to our results, AMGs had the strongest inhibitory effect on *E. tarda*, followed by *F. columnare* and *V. vulnificus*, but was less effective on *S. agalactiae* and *A. hydrophila*. Simora et al. (2021) conducted a bacterial killing kinetic assay to determine that cathelicidin and cecropin can reduce *F. columnare*/*E. tarda*/*A. hydrophila* counts in catfish kidney and liver. Regarding AMG-integrated fish, the number of *V. vulnificus* and *S. agalactiae* gradually reduced in epinecidin1-transgenic zebrafish (Peng et al., 2010). Moreover, TP3 or TP4 in tilapia muscle tissues efficiently decreased CFU after both *V. vulnificus* and *S. agalactiae* infections (Lin et al., 2016). In contrast, TH2-3 significantly inhibited the bacterial growth in transgenic zebrafish after *V. vulnificus* but not *S. agalactiae* infection (Hsieh et al., 2010).

These findings supported our meta-results in that epinecidin1, TP3, and TP4 was more effective in killing bacteria than TH2-3. What’s more, the reduction in CFU did not appear until 6 to 24 hours after pathogens have infected fish. Comparatively, immune-related gene expression was upregulated at two hours of infection, but CFU dropped only after four hours of gene up-regulation. It implies that pathogen invasion first triggers immune-related gene responses and subsequently suppresses bacterial growth.

### 4.3. Enhanced enzymatic activity contributes to disease-resistant enhancement

Another critical indicator or parameter associated with disease-resistant enhancement is the enzyme activity, which can strengthen immune and antioxidant systems (Biller and Takahashi, 2018), including lysozyme, superoxide dismutase, catalase and glutathione peroxidase (Abdel-Wahab et al., 2021; Rashidian et al., 2021; Wang et al., 2021). Meanwhile, increasing numbers of research have illuminated that AMPs can enhance lysozyme activity (Wang et al., 2022b) since fish lysozyme has been demonstrated to have lytic/phagocytic action against infectious microorganisms and to be crucial for innate immunity (Saurabh and Sahoo, 2008). In addition, previous literature exemplified that AMPs noticeably increased the cumulative survival rate to enhance disease resistance against a variety of pathogens via improving antioxidant capacity in Asian catfish (*Clarias batrachus*) (Kumari et al., 2003), zebrafish (Rashidian et al., 2021), Pengze crucian carp (*Carassius auratus* var. *Pengze*) (Wang et al., 2021) and Nile tilapia (Abdel-Wahab et al., 2021). These findings suggest that disease-resistant enhancement could be correlated with enhanced lysozyme activity and antioxidant capacity, which in turn contribute to the accelerated protection of fish against pathogen infections.

As a previous study summarized that AMPs as feed additives can improve disease resistance, as evidenced by increased antioxidant enzyme activity, lysozyme activity and upregulation of immune-related gene expression (Wang et al., 2022b). Generally, AMG-integrated fish encoding an AMP could confer a similar function by expressing mature AMPs once they are integrated into the genomes via genetic engineering. However, only the serum LYA was investigated in AMG-integrated fish to date. Mao et al. (2004) stated that the lysozyme activity did not increase significantly after *A. hydrophila* infection in lactoferrin-transgenic grass carp. Contrariwise, recombinant goose lysozyme in channel catfish exerted higher lysozyme activity after incubation with *M. lysodeikticus* (Pridgeon et al., 2013b). Collectively, our results showed that AMGs had a large effect on improvement of LYA, which were slightly different from previous individual studies. On the one hand, meta-analysis can increase sample size to eliminate heterogeneity between individual studies and obtain robust conclusions; nonetheless, the reliability of these results is highly dependent on the number of the studies involved. On the other hand, only three studies were included for LYA meta-analysis in the present study, and such a small sample size may draw conclusions that deviated from the actual results. Therefore, more related studies should be involved in the future analysis of the effect of AMG on enzymatic activity.

### 4.4. High cumulative survival rates denote enhanced resistance to pathogens

Although all recent investigations have demonstrated that the introduction of exogenous AMGs considerably boosts fish survival regardless of the type of bacteria and fish species employed, and a higher CSR than controls is directly related to improved disease resistance. There are still great differences in survival rates among various studies, including AMG-transgenic and control groups, and these significant variations contribute to a high heterogeneity (*I*^2^ = 97.25%). In this instance, our moderator-analysis denoted that the differences in AMGs, pathogenic type and fish species were responsible for varying CSR. In detail, AMGs played a greater role in improving the CSR of grass carp, medaka and zebrafish than in channel catfish and rainbow trout. With respect to the type of AMGs, cecropin, cecropin analog (CF-17) and cathelicidin were more powerful than other AMGs at enhancing CSR, and these AMGs exerted robust inhibitory effects against bacteria and viruses but not parasites.

Our meta-data suggested that the average CSR of the AMG-integrated individuals was three times higher than that of the control group (70.50% *vs* 21.43%). From the first AMG-transgenic catfish, Dunham et al. (2002) concluded that cecropin can enhance CSR from 27.30% to 95.78% when channel fish were infected with *F. columnare*, and a similar positive result was also determined in medaka (Sarmasik et al., 2002) Additionally, survival improvements of more than 30% were achieved in lactoferrin-transgenic grass carp, lysozyme-/hepcidin-transgenic zebrafish, cecropin-transgenic rainbow trout and cathelicidin-transgenic catfish against a variety of bacteria was achieved, respectively (Mao et al., 2004; Yazawa et al., 2006; Pan et al., 2011; Chiou et al., 2014; Wang et al., unpublished data). Regarding the viruses, grass carp hemorrhage virus (GCHV) and infectious hematopoietic necrosis virus (IHNV) were investigated in lactoferrin-transgenic grass carp and cecropin-/CF-17-transgenic rainbow trout, respectively. And previous studies demonstrated that survival increases of up to 40% and 60% were observed in in the respective two viruses. (Zhong et al., 2002; Chiou et al., 2014). However, Elaswad et al. (2019) reported that there was a slight increase without significant differences in CSR of cecropin-transgenic catfish compared to the wild-type group after the *Ichthyophthirius multifiliis* infection (55.75% *vs* 48.70%). Therefore, our meta-findings were supported by these individual studies.

### 4.5. Increased tolerance towards pathogens following engineered expression of AMGs

By integrating AMGs into the target species’ genome and expressing their encoded AMPs, AMG-genetic engineering aims to exert anti-infective effects on the target species. Therefore, the focus of exogenous AMG introduction should be to ensure that the gene can be expressed in the presence or absence of pathogen invasion. Accumulated evidence from recent studies has demonstrated that transgenic or genome editing technology can effectively improve disease resistance by integrating the modified constructs containing an AMG and promoter into the target fish.

Currently, the expression profiles of six AMGs were revealed in zebrafish and channel catfish. Interestingly, the expression of exogenous AMGs was maintained at a certain level even in the absence of pathogen invasion according to our results (Fig. 5A). What’s more, in contrast to the expression patterns of immune-related genes, our meta-analytic results determined that AMGs did not express in a tissue- or fish-specific manner. High expression of AMGs in the spleen, skin, muscle, liver and kidney of a variety of fish, as well as in certain other tissues, is encouraging and indicative of potential disease resistance in the current investigation. Pridgeon et al. (2013a) found that chicken-type lysozyme showed higher expression levels in the spleen and posterior kidney after *A. hydrophila* infection, but a lower level was represented in blood, skin and liver. Besides, the mRNA of GRN1 was abundantly expressed in the spleen, liver, muscle and kidney of GRN-41 transgenic zebrafish, followed by the skin, gill and head (Wu et al., 2018). A recent study determined that a weak expression of cathelicidin was detected in fin, intestine and head of the transgenic catfish without pathogen infections (Simora et al., 2020). This evidence suggested that the expression of exogenous AMG is not limited to some specific tissues and but is instead represented in different tissues at varying expression levels regardless of the type of pathogens, which is in agreement with our findings.

**Fig. 5.**
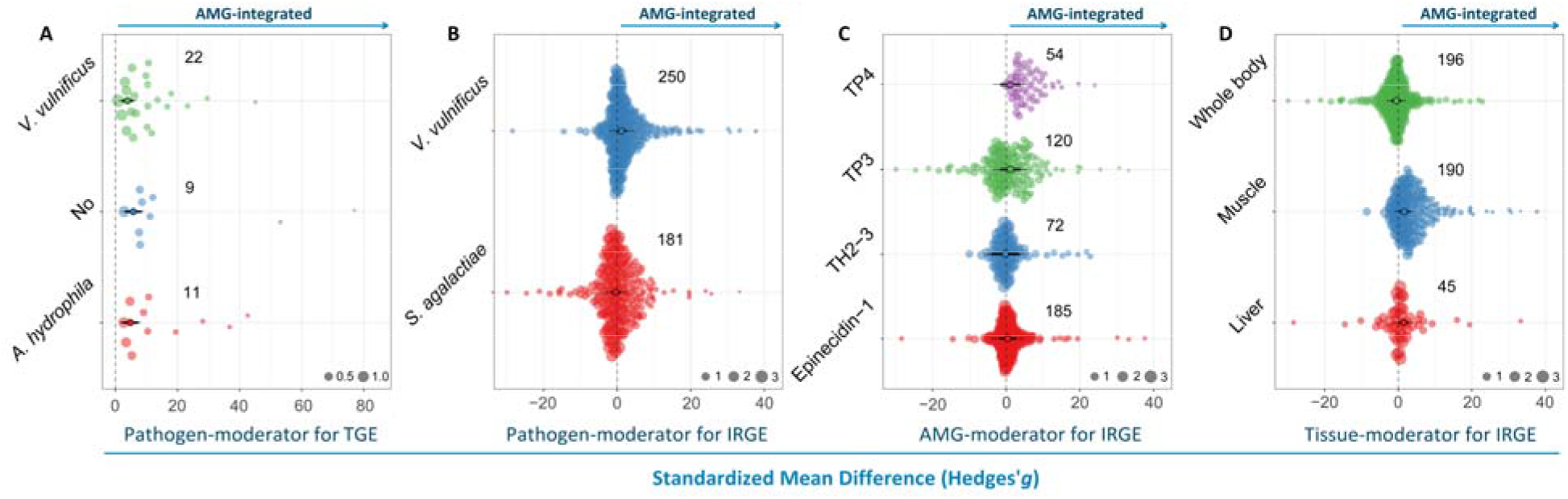
Results of moderator analysis after heterogeneity reduction. (**A**) The effect of pathogen type on the expression of AMGs (TGE) based on pathogen-moderator analysis. (**B**) The effect of pathogen type on the expression of immune-related genes (IRGE) using pathogen-moderator analysis. (**C**) The effect of AMG type on IRGE using AMG-moderator analysis. (**D**) The effect of tissue on IRGE using tissue-moderator analysis. AMG, antimicrobial peptide gene. For more thorough schematic details, please see Figure 2.

Indeed, the expression of AMG is often related to the type of promoter used, and tissue-specific promoters can limit the overexpression of AMG in specific tissues. For example, PGRN-transgenic zebrafish driven by MLC2 promoter exhibited a muscle-specific expression pattern of tilapia secreted PGRN peptides (Ju et al., 2003; Wu et al., 2018). Although PGRN-transgenic zebrafish expressed PGRN mRNA dominantly in muscle, abundant expressions were detected in immune-related organs, such as kidney, liver and spleen after *V. vulnificus* challenge (Wu et al., 2018). These findings imply that, even when a specific-expressed promoter is used for plasmid construction, the expression pattern of AMGs will not be easily altered, particularly after pathogen infections.

### 4.6. Synergies between AMGs and immune-related genes improve disease resistance

An increasing number of studies have unveiled that AMGs may exhibit multifaceted immunomodulatory properties via altering IRGE in various fish. Furthermore, the overexpression of exogenous AMGs is often accompanied by the body’s *in vivo* immune response, especially the coordinated expression of immune-related genes. Currently, exogenous AMGs have been demonstrated to be able to increase IRGE in grouper, Nile tilapia and zebrafish; however, the effect has only been observed in TP3-, TP4-, TH2-3-, and epinecidin1-transgenic fish, and most studies focused on *V. vulnificus* and *S. agalactiae*. Additionally, *V. vulnificus* infection or TP3/TP4 was more likely to induce the IRGE compared to other pathogens or AMGs (Fig. 5BC). In contrast to AMG expression, the IRGE typically requires a pathogen infection to be present. Our tissue-moderator analytic results confirmed that the distribution of IRGE is mainly distributed in liver and muscle (Fig. 5D). Notably, zebrafish are commonly used in transgenic models for improved disease resistance, and scientists typically extract RNA from the entire body rather than specific tissues due to their small size. Consequently, the profiles of IRGE in different tissues could not be determined even though our current findings can confirm that IRGE is presented in the zebrafish body.

In general, the patterns of IRGE changed over time, and IRGE was activated within two hours after fish infection, then increased, remained a high level at 4 to 12 hours, and subsequently reduced to the initial stage of infection at 12 to 48 hours. Besides, different genes tended to present different expression profiles over time even though they were induced by the same pathogen in one fish species. Emerging evidence supported that the expression of cytokines was elevated in epinecidin-transgenic zebrafish at various times after bacterial infections compared to non-edited fish, such as MyD88 at 2 hours, IL1β at 4 hours and IL10 at 8 hours post-infection (Peng et al., 2010). Inversely, the expressions of TNF and NFKβ were activated at 3 hours but returned to a low level at 48 hours after *V. vulnificus* infection (Lee et al., 2013). The variability in these expression patterns suggests that immune-related genes have feasible against a variety of pathogens. However, the temporal pattern of the synergistic expression of AMGs and immune-related genes cannot be accurately predicted due to the lack of time-varying expression of AMGs, even though the expression trends of immune-related genes over time have been determined.

## 5. Conclusions and perspectives

Meta-analysis has developed into a potent and popular method for integrating data from multiple studies and guiding scientifical decision-making in the life sciences community. The results of our integrative analysis revealed that AMG-integrated fish exhibited broad-spectrum antibacterial properties and a higher cumulative survival rate after pathogen invasion compared to non-AMG-edited individuals. Furthermore, this strong evidence confirms the feasibility of AMGs to accelerate the improvement of disease resistance by harnessing genetic engineering. Nonetheless, the combat against pathogens is protracted, and new approaches and strategies should be applied to broaden the scope of AMGs as alternatives to conventional antibiotics.

Genome editing dominated by CRISPR-based platforms has risen rapidly in life sciences over the past decade. In aquaculture, the CRISPR/Cas9-mediated system holds a promise for the enhancement of favorable traits, especially growth, disease resistance, sterility and fatty acid. In addition to being affordable and effective, CRISPR/Cas9 has the property of being widely applicable because it allows for simultaneous modifications to various sites via delivering multiple sgRNAs with the Cas9 protein/mRNA (Yang et al., 2013; Ota et al., 2014). As previously established, AMGs can successfully improve fish disease resistance through transgenic integration. However, the maximum of disease-resistant enhancement has been somewhat constrained by the fact that the majority of these vast research have thus far concentrated on a single AMG induction. Theoretically, it is conceivable to induce two or more AMGs into the genome to acquire hereditary higher resistance to diseases by using CRISPR/Cas9 genome editing compared to one AMG.

In addition to integrating exogenous AMGs into the genome, great achievements have demonstrated that knockout of the immune-related genes can enhance resistance against pathogens through negatively regulating gene expression or disrupting pathways. For instance, a few representative genes, like rhamnose-binding lectin (RBL), STAT2, JAM-A and PoMaf1, have undergone mutations that mimicked and altered the immunity of the fish and improved the host’s resistance to disease (Wang et al., 2022a). Indeed, RBL is a critical component of fish’s innate immunity as an antibacterial and non-self-recognition molecule (Booy et al., 2005; Watanabe et al., 2009), especially in the protection of teleost eggs as well as in the mucosa (Beck et al., 2012). Interestingly, a potential negative regulation of RBL is involved in the immunity of some fish against pathogenic invasion. Beck et al. (2012) confirmed that columnaris susceptibility was negatively linked with RBL expression levels. Furthermore, vulnerable fish’s gills showed higher up-regulated levels of RBL than those of resistant fish (Peatman et al., 2013). Recently, an RBL-mutated channel catfish line was established (Elaswad et al., 2018), and a higher survival rate was represented in their F_2_ individuals compared to those wild-type fish after being infected with *Flavobacterium columnare* (unpublished data). In this regard, integrating AMGs into these susceptibility loci can bidirectionally boost disease resistance based on gene pleiotropy. Alternatively, some studies have proved that myostatin(MSTN)-deficient fish not only grow faster, but also reduce disease susceptibility to *Edwardsiella ictaluri* in channel catfish (Coogan et al., 2022). Therefore, MSTN is also an alternative locus of an AMG integration for enhanced disease resistance.

With respect to sterilized or fatty acid-enriched fish lines, similar strategies were adopted. The hypothalamus-pituitary-gonad (HPG) axis is primarily responsible for controlling gonadal development and maturation in fish, and gonadotropin-releasing hormone (GnRH) is essential for regulating the differentiation of the gonad through the HPG axis (Mylonas et al., 2010). Additionally, follicle-stimulating hormone (FSH) and luteinizing hormone (LH) play different roles in early and late reproductive cycles, respectively, but they are both regulated by GnRH (Ogiwara et al., 2013). Genes encoding these three key hormones are pivotal in gonadal maturation. Qin et al. (2016) and Qin et al. (2022) revealed that channel catfish with LH- or GnRH-mutations had a reduced fertility compared to wild-type individuals, indicating that knocking out or disrupting the LH or GnRH gene can induce reproductive confinement. Docosahexaenoic acid (DHA), an omega-3 fatty acid, is essential for the development of human eye and nerve tissues (Judge et al., 2007), and the DHA biosynthesis pathway requires the elongase gene Elovl2 to function (Gregory and James, 2014). Recent studies showed that transgenic fish carrying elongase-like or Elovl2 gene exhibited higher DHA content compared to non-edited individuals (Alimuddin et al. 2008; Xing et al., 2022). Thus, compared with knocking out reproductive-related genes to achieve sterilization, DHA-enrich fish are usually created by knocking in elongase-like genes.

Beyond the successful achievements of one desired trait, it is possible to use CRISPR/Cas9 to simultaneously improve multiple characteristics based on these empirical data and theoretical foundations, which means that we could alter other traits through the construction of different vectors while we focus on disease-resistant enhancement. From a genetic perspective, our hypothesis is that replacing the original functional genes with AMGs in specific coding regions of the chromosome would confer multi-generational antimicrobial activities of the host and momentously improve multi-valuable traits. This strategy will hopefully allow us to create new fish lines possessing multiple favorable traits, such as sterilized and disease-resistant, growth-boosted and disease-resistant, or DHA-enriched and disease-resistant, or hybrid lines that have all of these traits (Fig. 6). In this vein, an example of our team demonstrates that it is highly feasible to insert the cathelicidin gene at the LH locus and cecropin gene at MSTN locus using a one-step CRISPR/Cas9-mediated system, resulting in gene-edited fish with increased disease resistance and growth but decreased fecundity (unpublished data). However, site-directed knock-in of multi-locus genes tends to increase mosaicism and off-target (Hsu et al., 2013; Yang et al., 2013). In this scenario, we should combine genetic engineering with selective breeding to maximize transgenic performance, avoiding malformations or unintended phenotypes. That is, the available gene-edited individuals with target traits are selected first, and homozygotes are produced by crossing breeding, which can more effectively integrate multiple desired traits into one individual. These theoretically executable hypotheses should be validated by reliable experimental evidence in future gene editing designs.

**Fig. 6.**
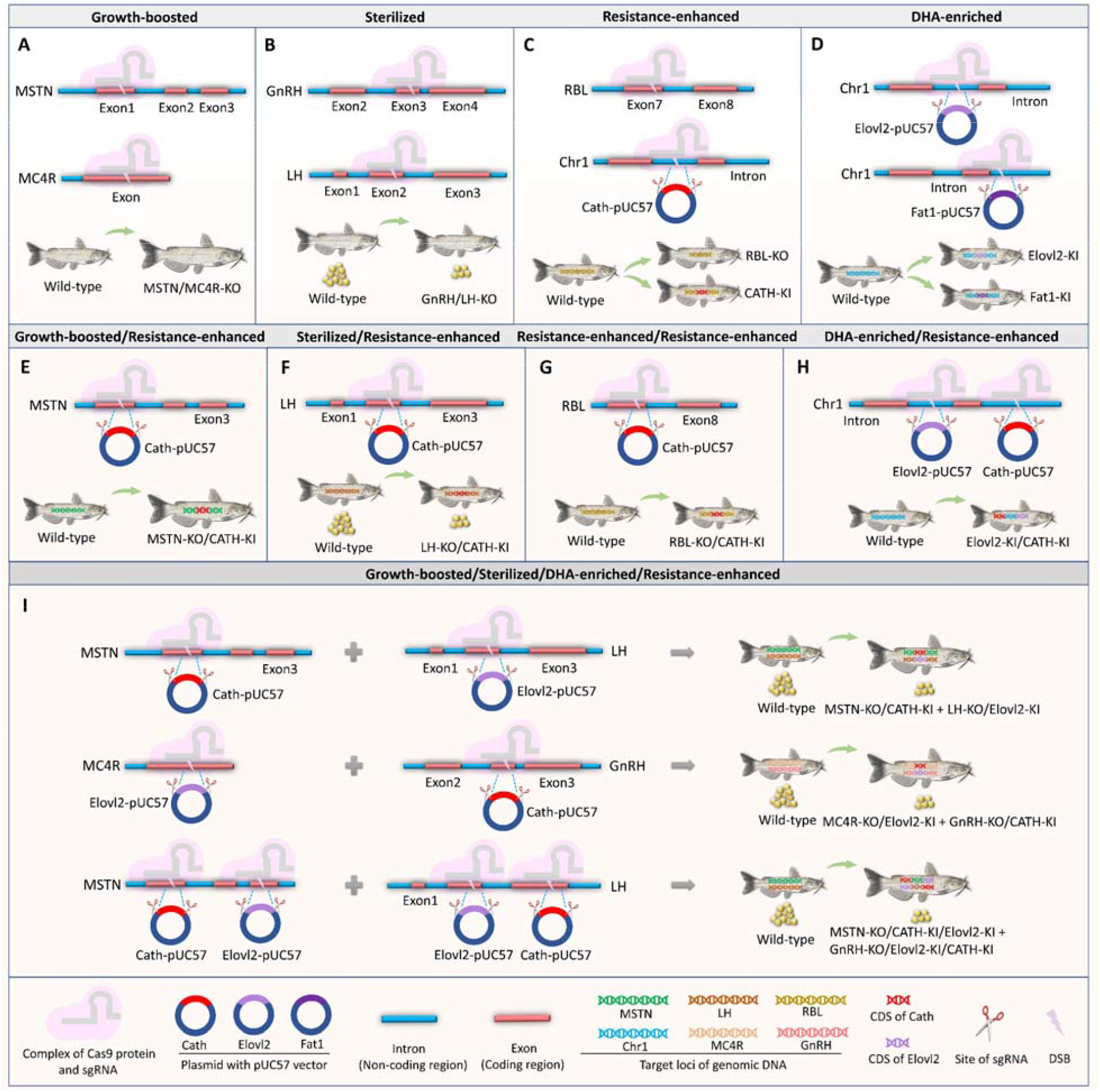
CRISPR/Cas9-mediated system induces traits of interest to disease resistance combined with sterilized, growth-boosted and DHA-enriched characteristics. (**A**) Growth-boosted fish line was created through knocking out MSTN or MC4R gene. (**B**) Sterilized fish line was produced through knocking out GnRH or LH gene. (**C**) Resistance-enhanced fish line was created through knocking out RBL or knocking in Cath gene at the non-coding region of chromosome 1. (**D**) DHA-enriched fish line was generated through knocking in Elovl2 or Fat1 gene at the non-coding region of chromosome 1. (**E**) Resistance-enhanced fish with fast-growing was produced by knocking in Cath gene at the MSTN locus. (**F**) Resistance-enhanced fish with sterility was produced by knocking in Cath gene at the LH locus. (**G**) A higher resistance-enhanced fish line was created by knocking in Cath gene at the RBL locus. (**H**) Resistance-enhanced fish with high DHA content was produced by knocking in Cath and Elovl2 genes at the non-coding region of chromosome 1. (**I**) Multiple CRISPR/Cas9 systems produce hybrid fish lines that contain enhanced-resistance, fast-growing, sterility and enriched-DHA traits. MSTN, myostatin; MC4R, melanocortin 4 receptor; GnRH, gonadotropin-releasing hormone; LH, luteinizing hormone; Chr1, chromosome 1; KO, knock out; KI, knock in; Cath-/Elovl2-/Fat1-pUC57, a plasmid containing cathelicidin/elovl2/fat1 gene constructed with pUC57 as the vector; CDS, coding sequences; DSB, double-stranded break.

## Data availability

Supplementary data to this article can be found online.

## Acknowledgements

This work was supported by the China Scholarship Council, grant number (CSC201906330109 and CSC201908160003).

